# Adapting the endoplasmic reticulum proteostasis rescues epilepsy-associated NMDA receptor variants

**DOI:** 10.1101/2023.04.01.535233

**Authors:** Pei-Pei Zhang, Taylor M. Benske, James C. Paton, Adrienne W. Paton, Ting-Wei Mu, Ya-Juan Wang

## Abstract

The *GRIN* genes encoding N-methyl-D-aspartate receptor (NMDAR) subunits are remarkably intolerant to variation. Many pathogenic NMDAR variants result in their protein misfolding, inefficient assembly, reduced surface expression, and impaired functionality at the plasma membrane, causing neurological disorders including epilepsy and intellectual disability. Here, we concentrate on the proteostasis maintenance of NMDARs containing epilepsy-associated variations in the GluN2A (or NR2A) subunit, including M705V and A727T. We showed that these two variants are targeted to the proteasome for degradation and have reduced functional surface expression. We demonstrated that the application of BIX, a known small molecule activator of an HSP70 family chaperone BiP (<u>B</u>inding immunoglobulin <u>P</u>rotein) in the endoplasmic reticulum (ER), significantly increases total and surface protein levels, and thus the function of the M705V and A727T variants in HEK293T cells. Mechanistic studies revealed that BIX promotes folding, inhibits degradation, and enhances anterograde trafficking of the M705V variant by modest activation of the IRE1 pathway of the unfolded protein response. Our results showed that adapting the ER proteostasis network restores the folding, trafficking, and function of pathogenic NMDAR variants, representing a potential treatment for neurological disorders resulting from NMDAR dysfunction.

## Introduction

N-methyl-D-aspartate receptors (NMDARs) are excitatory neurotransmitter-gated ion channels in the mammalian central nervous system and play a key role in mediating synaptic plasticity and maintaining the excitation-inhibition balance within synapses [1, 2]. NMDARs are tetramers that are assembled from two obligatory GluN1 subunits and two GluN2 (A to D) subunits or two GluN3 (A and B) subunits. Each of these subunits shares a common domain architecture, including an extracellular amino-terminal domain (ATD), an extracellular ligand-binding domain (LBD), a transmembrane domain comprised of three transmembrane helices and one reentrant loop, and an intracellular carboxy-terminal domain (CTD) (**Fig. 1a**) [3, 4]. The most common subtype in the human brain is constructed from two GluN1 (or NR1) subunits and two GluN2A/2B (or NR2A/2B) subunits [5–7]. NR1NR2 channels are activated by the simultaneous binding of an agonist (glutamate or NMDA) in the NR2 subunits and a co-agonist (glycine or d-serine) in the NR1 subunits to the LBD. Numerous pathogenic variants in NR2A subunit-encoding gene, *GRIN2A*, have been identified in patients with a variety of neurological and neurodevelopmental disorders including epilepsies, autism spectrum disorder, and intellectual disabilities, with patients displaying phenotypes of cognitive dysfunction, dyspraxia, developmental delay, and speech difficulties [8, 9]. These NR2A variants lead to impaired signaling at the plasma membrane [10]. Although recent advances in whole genome sequencing have identified an increasing number of pathogenic variants in various subunits of NMDARs, the precise mechanism of how these variants influence receptor surface delivery and function remains poorly understood [11].

**Figure 1.**
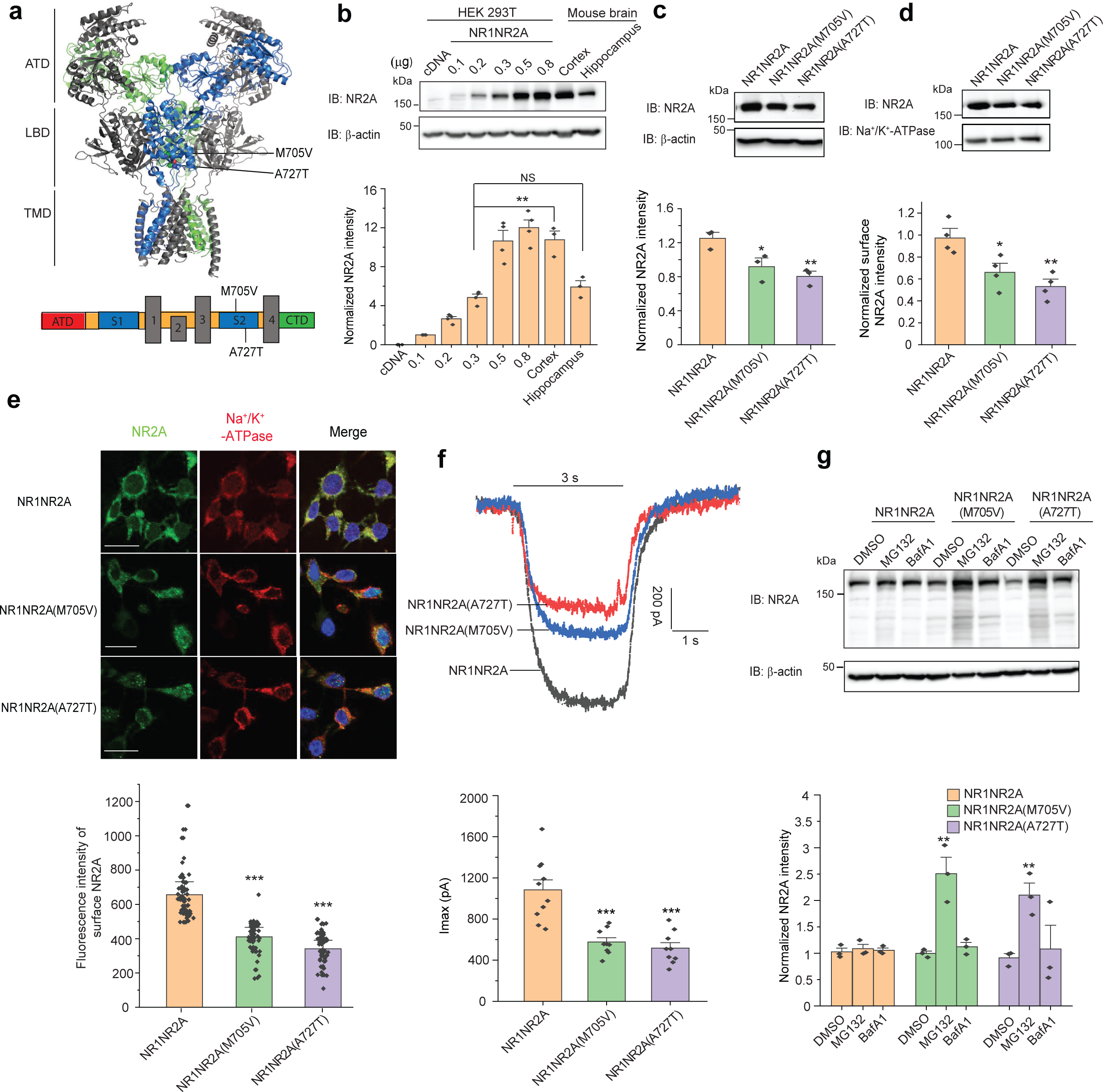
**The alternations as a result of variants in NR2A subunit on the expression and function of trafficking-deficient NMDARs. a**, Upper panel, cartoon representation of tetrameric NMDARs, with the NR1 subunits in light and dark grey and the NR2A subunits in green and blue (PDB: 7EOS [52]), each subunit consisting of an extracellular amino-terminal domain (ATD), an extracellular ligand-binding domain (LBD), a transmembrane domain comprised of three transmembrane helices and one reentrant loop (TMD), and an intracellular carboxy-terminal domain (CTD). The positions of the M705V and A727T are highlighted as spheres in LBD. Lower panel, linear representation of an NR2A subunit. The LBD contains two segments of S1 and S2. **b**, The NR2A expression level in transiently transfected HEK293T cells compared to the cortex and hippocampus tissue in mouse brain. NS:not significant. **c**, The effect of M705V and A727T on the total protein expression level of NR2A. β-actin serves as the loading control of total protein lysates. **d**, The effect of M705V and A727T on the surface protein expression level of NR2A as determined by surface biotinylation assays. Na^+^/K^+^-ATPase serves as the loading control of membrane proteins. **e**, Surface NR2A staining was in green (column 1), and plasma membrane marker Na^+^/K^+^-ATPase staining was in red (column 2). Merge of these two signals with the nucleus stained in blue with DAPI was shown in column 3. Scale bar = 20 μm. The fluorescence intensity of the surface NR2A was quantified from 30-50 cells per condition as shown on the lower panel. **f**, Automated patch-clamping was performed with the IonFlux Mercury 16 ensemble recording at a holding potential of -60 mV. Glutamate (10 mM) and glycine (100 µM) were applied simultaneously for 3 s, as indicated by the horizontal bar above the currents. The peak currents (Imax) were acquired and analyzed by the Fluxion Data Analyzer (n = 9 - 10 ensemble recording; each ensemble recording enclosed 20 cells). **g**, Inhibition of proteasomal degradation (MG132, 10 µM, 6 h) and lysosomal degradation (BafA1, 1 µM, 6 h) independently demonstrated that NR2A subunits containing either M705V or A727T variants are mainly degraded by the proteasome pathway. Each data point is reported as mean ± SEM. One-way ANOVA followed by a post-hoc Tukey test was used for statistical analysis. *, p < 0.05; **, p < 0.01; ***, p < 0.001.

We focus on proteostasis maintenance of neurotransmitter-gated ion channels, including NMDARs [11, 12]. NMDAR subunits are folded and assembled to heterotetramers in the endoplasmic reticulum (ER), a cellular organelle for protein quality control with the assistance of molecular chaperones. The assembled receptors engage the trafficking machinery to exit the ER and travel through the Golgi apparatus en route to the plasma membrane. Reduced surface expression of variant NMDARs is a major molecular mechanism in the pathogenesis of related neurological diseases, such as epilepsy and intellectual disability [1, 8, 11]. Accumulating evidence indicates that the structural integrity of the LBD is essential for the surface trafficking of NMDARs [13–15]. Therefore, many disease-associated variations in the NR2A subunit that are located in the LBD cause the reduced presence of the NMDARs on the plasma membrane and thus loss of their function and epilepsy syndrome [8, 11]. For example, two missense variations, M705V and A727T in the S2 pocket of the LBD of NR2A subunit (**Fig. 1a**), which were identified in a cohort of patients with epilepsy, impair the delivery of the mature NMDARs to the plasma membrane [13, 16]. Presumably, trafficking-deficient variants in NMDAR subunits result in their protein misfolding and retention within the ER, which leads to their excessive degradation via the ER-associated degradation (ERAD) pathway [17]. Consequently, this can contribute to the loss-of-function phenotype displayed by variant NMDARs and corresponding disease phenotypes due to the inefficient processes of folding, assembly, and trafficking.

There is limited knowledge in the literature about how the proteostasis network regulates the folding, assembly, degradation, and trafficking of NMDARs [11]. The ER proteostasis network regulates the ER folding capacity to assure that newly synthesized proteins achieve their native multidimensional structures in the oxidative folding environments [18, 19]. Since membrane proteins need to fold in the ER, pharmacologic enhancement of ER proteostasis capacity to restore variant-containing NMDAR folding and assembly in the ER has the potential to improve the anterograde trafficking and function of NMDARs at the cell surface [20–22]. BiP (Binding immunoglobulin Protein, Grp78), an ER resident HSP70 family chaperone, plays a prominent role in the ER by controlling protein folding and preventing aggregation [23]. BiP is the master regulator of the unfolded protein response (UPR), which monitors the ER proteostasis status [24]. Activation of the UPR is a promising way to change the fate of pathogenic proteins that are associated with various protein misfolding diseases [22, 25, 26]. The UPR contains three integrated signaling arms, including ATF6 (Activating Transcription Factor 6), IRE1 (Inositol-Requiring Enzyme 1), and PERK (Protein Kinase R-like ER Kinase) [24, 25]. In response to ER stress, these activated pathways result in both translational and transcriptional signaling cascades that function to regulate diverse aspects of cellular physiology including ER proteostasis. However, it remains unclear whether the UPR activation restores proteostasis of pathogenic, misfolded NMDAR proteins harboring clinical variants.

Here, we applied BIX (1-(3,4-dihydroxy-phenyl)-2-thiocyanate-ethanone), a potent BiP activator, [27, 28] to investigate how regulating the ER proteostasis network influences the folding, trafficking, and function of NMDARs carrying M705V and A727T variants in the LBD of the NR2A subunit.

## Results

### Epilepsy-associated variants in the NR2A subunits reduce the protein expression and function of NMDARs

To evaluate the protein levels of exogenously expressed NMDARs in HEK293T cells compared to those in the central nervous system, we transiently transfected HEK293T cells with different amounts of wild type (WT) NR1 and NR2A plasmids in a 6-well plate and compared NR2A expression to endogenous NR2A protein levels in the adult mouse cortex and hippocampus (**Fig. 1b**). Mouse cortex and hippocampus demonstrated high expression levels of NR2A, which was consistent to previous reports [29–31]. We observed that transient transfection using 0.3 μg NR1 and 0.3 μg NR2A plasmids per well led to comparable NR2A protein level as expressed in mouse hippocampus. Therefore, in the following experiments, we kept this transfection regime to express low levels of exogenous NMDARs in HEK293T cells to minimize potential disturbance to protein homeostasis, yet to achieve appreciable NR2A protein levels that are physiologically relevant. Additionally, to lower the excitatory toxicity that associated with the expression of NMDA receptors, cells are routinely maintained in cell culture media supplemented with 50 µM of MK801 and 2.5 mM of MgCl2 to block the channels [32].

We determined the effect of the M705V and A727T variants, located in the S2 segment of the LBD of the NR2A subunit (**Fig. 1a**), on the trafficking and function of NMDARs [33, 34]. Western blot analysis of cell lysates from the transfected HEK293T cells showed that NR2A total protein levels were significantly reduced for the M705V and A727T variants (**Fig. 1c**). Because NMDARs need to be transported to cell surface to function as ion channels, we determined whether variant NMDARs affected the cell surface expression. Surface biotinylation assay demonstrated that the surface protein level was significantly reduced in the NR2A subunits containing M705V or A727T in HEK293T cells (**Fig. 1d**). In addition, we used immunocytochemistry to visualize surface expression of NR2A subunits. Cells were fixed with 4% formaldehyde without membrane permeabilization and incubated with an anti-NR2A antibody that has an epitope recognizing the NR2A extracellular region. NR2A surface staining merged well with the staining of a plasma membrane marker Na^+^/K^+^ ATPase (**Fig. 1e**). The fluorescence intensity of surface NR2A was calculated for WT and each variant [35]. The immunocytochemistry analysis showed that M705V or A727T significantly reduced the NR2A cell surface expression in comparison to WT (**Fig. 1e**), indicating that the variants have impaired efficiency to traffic to the plasma membrane.

Moreover, to determine whether NR2A variants influence the function of NMDARs, we carried out whole-cell patch-clamping electrophysiological recording in HEK293T cells using the Fluxion automated patch clamping instrument. Application of co-agonists of 10 mM glutamate and 100 µM glycine opened the NMDAR channels and showed the slowly decaying currents (**Fig. 1f**), consistent with the electrophysiological characteristics of the channels [1]. The peak agonist-induced current in the WT NR1NR2A group was 1083.9 ± 305.2 pA, whereas the peak current in NMDARs harboring NR2A(M705V) or NR2A(A727T) decreased to 576.9 ± 122.7 pA and 517.4 ± 156.5 pA, corresponding to 46.8% and 52.3% reduction, respectively (**Fig. 1f**), indicating the loss-of-function phenotype of these NR2A variants.

Collectively, we demonstrated that the M705V and A7275T variations significantly reduced the steady-state protein levels and the surface expressions of NR2A and peak current amplitudes of NMDARs. Ion channel mutations often cause reduced protein levels due to the degradation of misfolded proteins by the protein quality control machinery. Misfolded proteins can be targeted to the proteasome or the lysosome for their destruction. Therefore, we investigated the preferred degradation route of the NR2A variants by inhibiting potential degradation pathways pharmacologically. Application of MG132 (10 μM, 6 h), a potent proteasome inhibitor, substantially accumulated both NR2A variants of M705V and A727T in cells according to western blot analysis without an apparent effect on the WT NR2A; in sharp contrast, application of bafilomycin A1 (Baf A1, 1 μM, 6 h), a potent lysosome inhibitor, did not change the total protein levels of NR2A(M705V) and NR2A(A727T) variants (**Fig. 1g**), indicating that these two variants are targeted to the proteasome for their degradation, presumably through the ERAD pathway. Since methionine can form stabilizing interactions with adjacent aromatic residues [36] and M705 is surrounded by F682, Y704 and F708 in NR1A (**Fig. S1a**), the M705V variation potentially destabilizes this protein motif, leading to its misfolding and degradation. Since A727 is located in the hydrophobic pocket composed of V535, V537, F682, and I729 (**Fig. S1b**), the A727T variation potentially compromises the hydrophobic interactions as well as introducing steric clashes, causing its excessive degradation.

### BIX increases functional surface protein levels of loss-of-function NR2A variants

We aim to correct the folding and surface trafficking of NR2A variants to restore their function on the plasma membrane. Previously it has been shown that BIX has a neuro-protective effect against ER stress [28] and we used BIX to restore the function of pathogenic neuroreceptors [37]. Since BIX has the capacity to enhance the ER folding capacity [20], we determined the effect of BIX application on variant NR2A protein levels in HEK293T cells stably expressing NR1NR2A(M705V) and NR1NR2A(A727T). Dose-response analysis showed that BIX (24 h treatment) increased total protein levels of NR2A variants in a dose-dependent manner: the EC50 (half-maximal effective concentration) values of BIX were 5.28 μM for NR2A(M705V) and 6.90 μM for NR2A(A727T) (**Fig. 2a**). BIX increased NR2A variant protein levels at a concentration above 2.5 μM and the effect of BIX plateaued at a concentration of 10 μM (**Fig. 2a**). Resazurin cell toxicity assay showed that a single dose application of BIX for 24 h did not significantly induce toxicity to cells at concentrations below 10 μM (**Fig. 2b**). Time-course experiments demonstrated that the increase of total NR2A(M705V) or NR2A(A727T) subunit protein levels by a single dose application of BIX (10 μM) was achieved as early as 6 h and maximized at 12- 24 h after treatment (**Fig. 2c**). Thereby, we used the optimal treatment condition of BIX (10 μM, 24 h) for the following experiments. BIX treatment did not change the total protein levels of wild type NR2A subunit in human A2780 cells that express endogenous NMDA receptors (**Fig. S2**) [38], indicating that BIX can achieve certain selectivity toward the destabilized ion channel variants.

**Figure 2.**
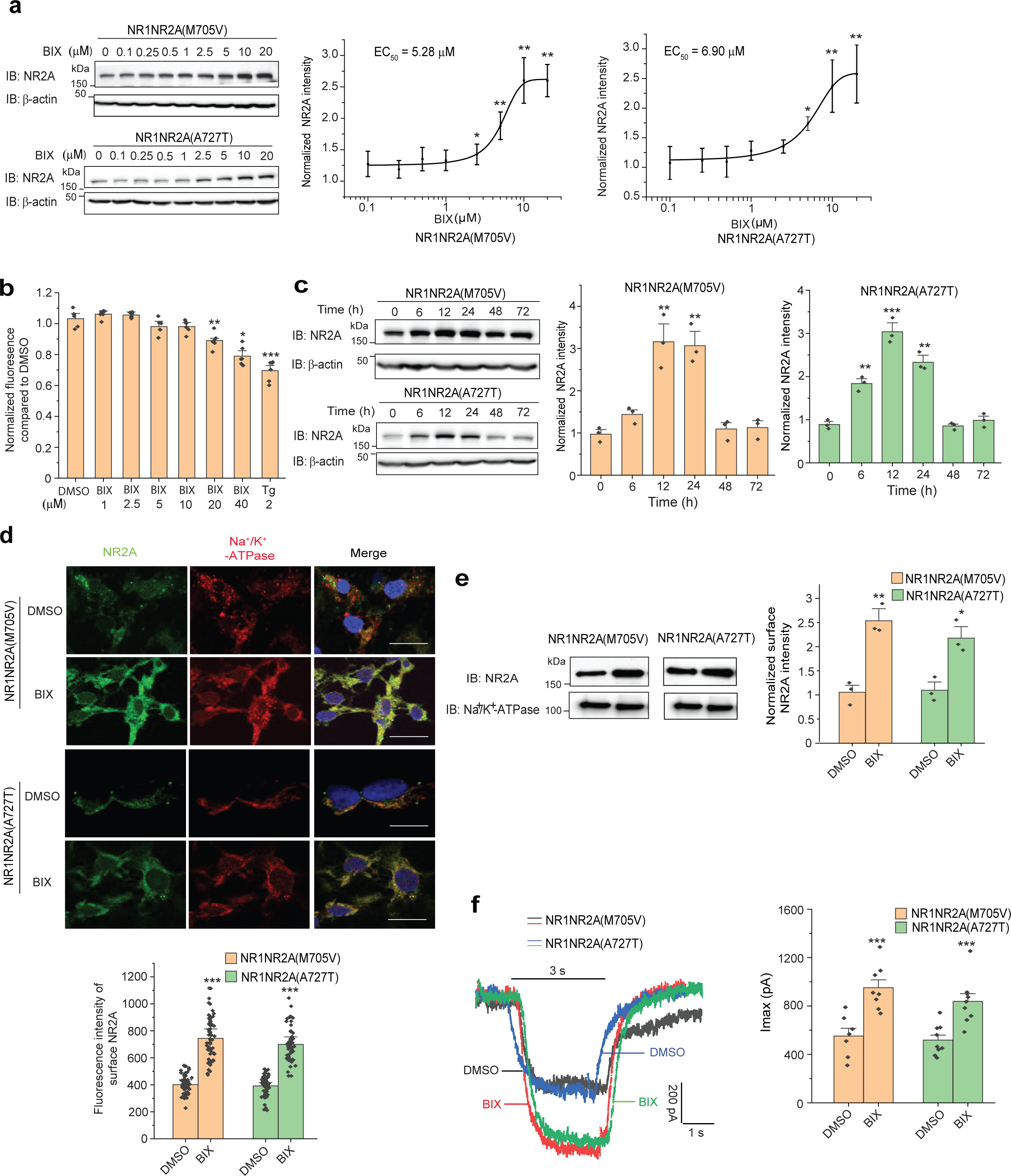
**Effects of BIX on the expression and function of trafficking-deficient variants in NR2A subunit of NMDARs. a**, Dose-response analysis of BIX treatment (24 h) on the total protein levels of NR2A subunits in HEK293T cells expressing NR1NR2A(M705V) or NR1NR2A(A727T). **b**, The toxicity of BIX treatment (24h) was quantified with a resazurin assay to determine cell viability under different concentrations of BIX application. Tg (2 µM, 6 h) was used as a positive control. Tg: thapsigargin. **c**, Time-course analysis of BIX (10 μM) treatment on the total protein levels of NR2A subunits in HEK293T cells expressing NR1NR2A(M705V) or NR1NR2A(A727T). β-actin serves as the total protein loading control. **d,** BIX treatment (10 μM, 24 h) increases surface expression of variant NMDARs. Surface NR2A staining was in green (column 1), and plasma membrane marker Na^+^/K^+^-ATPase staining was in red (column 2). Merge of these two signals with the nucleus stained in blue with DAPI was shown in column 3. Scale bar = 20 μm. The fluorescence intensity of the surface NR2A was quantified from 30-50 cells per condition. **e**, Surface biotinylation assays further demonstrated that BIX (10 µM, 24 h) enhances surface expression of variant NMDARs. Na^+^/K^+^-ATPase serves as the loading control of membrane protein. **f**, BIX (10 μM, 24 h) restores function of variant NMDARs as ion channels, as shown by whole-cell patch-clamping recordings. Glutamate (10 mM) and glycine (100 µM) were applied simultaneously for 3 s, as indicated by the horizontal bar above the currents. The peak currents (Imax) were acquired and analyzed by the Fluxion Data Analyzer (n = 7 - 9 ensemble recording; each ensemble recording enclosed 20 cells). Student’s t-test (for comparison of two groups) or one-way ANOVA followed by a post- hoc Tukey test (for comparison of three or more groups) was used for statistical analysis. Each data point is presented as mean ± SEM *, p< 0.05; **, p< 0.01, ***, p < 0.001.

Since NMDARs need to reach the plasma membrane to carry out their function, we performed experiments to determine how BIX treatment affected the surface levels of NR2A variants. Immunofluorescence microscopy experiments demonstrated that BIX treatment substantially increased the surface staining of NR2A(M705V) and NR2A(A727T) subunits in HEK293T cells (**Fig. 2d**, column 1), which colocalized well with a plasma membrane marker Na^+^/K^+^ ATPase (**Fig. 2d**, column 3). Consistently, surface biotinylation assays revealed that BIX treatment significantly increased the surface protein levels of NR2A(M705V) and NR2A(A727T) as well (**Fig. 2e**).

To determine whether the enhanced surface trafficking of NR2A variants led to enhanced function of NMDARs, we carried out whole-cell patch-clamping experiments to record the agonist-induced peak currents of NMDARs. BIX treatment (10 μM, 24 h) significantly increased the peak current amplitudes from 552.1 ± 169.9 pA to 951.8 ± 183.0 pA for the NR1NR2A(M705V) variant, and from 518.4 ± 125.0 pA to 839.0 ± 190.1 pA for the NR1NR2A(A727T) variant, corresponding to 72.3% and 61.8% increase, respectively (**Fig. 2f**). These results unambiguously demonstrated BIX’s capability to upregulate the surface trafficking and the function of NMDARs containing the NR2A variants of M705V and A727T.

### BIX enhances variant NMDARs proteostasis by promoting their folding and trafficking and reducing their ERAD

Thereafter, we focused on the NR2A(M705V) variant to study the mechanism of action of BIX on restoring its surface trafficking and function. We determined whether BIX treatment enhanced folding and facilitated the anterograde trafficking of NR2A(M705V). To quantify the relative folding extent of NR2A(M705V), we used the n-dodecyl-β-d-maltoside (DDM) detergent solubility assay by measuring the ratio of its DDM-soluble / insoluble fractions in HEK293T cells. Application of BIX substantially increased this ratio for the variant NR2A(M705V) subunits (**Fig. 3a**), which indicated that BIX redistributed variant NR2A proteins from an aggregation-prone state to a pro-folding state.

**Figure 3.**
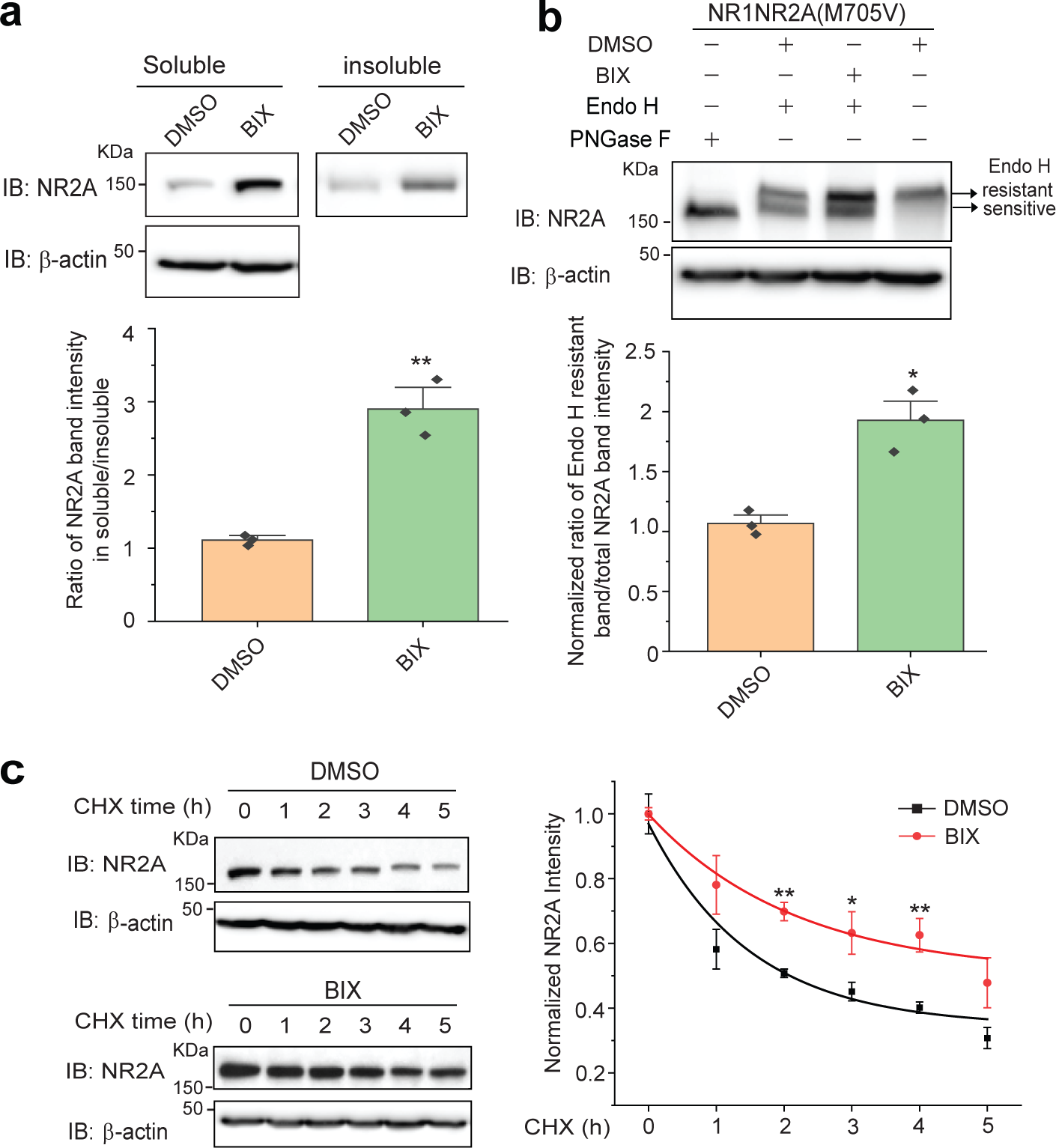
**Effect of BIX treatment on the folding, trafficking, and degradation of variant NMDARs. a**, Effect of BIX (10 μM, 24 h) on the folding of NR2A subunits in HEK293T cells stably expressing NR1NR2A(M705V). The normalized ratio of the soluble to insoluble NR2A was shown on the bottom panel. **b**, BIX (10 μM, 24 h) increases the Endo H-resistant post-ER glycoform of the NR2A subunit in HEK293T cells stably expressing NR1NR2A(M705V). PNGase F, which cleaves all glycans in a glycoprotein, served as a non-glycosylated form of NR2A. Quantification of the Endo H resistant/total NR2A band intensities was shown on the bottom panel. **c**, Effect of BIX (10 μM, 24 h) on the degradation of the NR2A subunit in HEK293T cells stably expressing NR1NR2A(M705V) using cycloheximide (CHX)-chase analysis. Each data point was reported as mean ± SEM. Student’s t-test (for comparison of two groups) or two-way ANOVA followed by a post-hoc Tukey test (for comparison of three or more groups) was used for statistical analysis. *, p < 0.05; **, p < 0.01.

In order to evaluate whether BIX enhanced the effective anterograde ER-to-Golgi trafficking of NR2A(M705V) subunits, we performed an endoglycosidase H (endo H) enzyme digestion experiment to quantify the trafficking efficiency. Endo H-sensitive NR2A subunits (**Fig. 3b**, lanes 2 and 3, bottom bands) are retained in the ER, either as nascent proteins or targeted for degradation, whereas Endo H-resistant NR2A subunits (**Fig. 3b**, lanes 2 and 3, top bands) have exited the ER and traffic at least to the Golgi for further glycan modifications. PNGase F treatment, which cleaves all glycans in a glycoprotein, generated a non-glycosylated form of NR2A (**Fig. 3b**, lane 1). Trafficking efficiency was quantified by the ratio of Endo H resistant NR2A / total NR2A bands. The group of BIX treatment produced a stronger post-ER endo H-resistant NR2A subunit band (**Fig. 3b**, **cf**. lane 3 to 2) and consistently increased the ratio of the endo H-resistant band / total NR2A, indicating that BIX significantly enhanced the trafficking efficiency of the variant NR2A(M705V).

Next, we determined whether BIX influenced the degradation of NR2A(M705V) variant. Cycloheximide, a potent protein synthesis inhibitor, was administered to the cell culture media for the indicated time. The cycloheximide-chase assay showed that BIX treatment significantly increased the normalized remaining NR2A(M705V) protein levels at 2h, 3h, and 4h after cycloheximide treatment (**Fig. 3c**), indicating that BIX attenuated the excessive degradation of variant NR2A(M705V). An exponential decay fitting showed that BIX treatment increased the half-life of NR2A(M705V) from 1.2 h to 2.7 h (**Fig. 3c**), indicating that BIX treatment reduced the degradation rate of this variant. Collectively, these data demonstrated that BIX enhanced the productive folding of the NR2A(M705V) variant in the ER, inhibited its ERAD, and promoted its anterograde trafficking from the ER to the Golgi and onward to the plasma membrane. Consequently, BIX treatment restored the function of NMDARs containing this variant due to the increased surface expression.

### BIX activates the IRE1 arm of UPR to increase the protein levels of variant NR2A(M705V) subunits of NMDARs

Furthermore, since the ER proteostasis network plays a critical role in regulating the biogenesis of ion channels [11, 12], we determined the effect of BIX treatment on this network. We investigated how BIX treatment influenced major ER chaperones [39, 40], including BiP, GRP94 (Glucose-Regulated Protein 94), calreticulin (CRT), and calnexin (CANX). Western blot analysis demonstrated that BIX treatment significantly increased the BiP protein level without an apparent effect on other selected chaperones in HEK293T cells expressing NR1NR2A(M705V) (**Fig. 4a**), consistent with the literature report that BIX is a potent BiP inducer [28].

**Figure 4.**
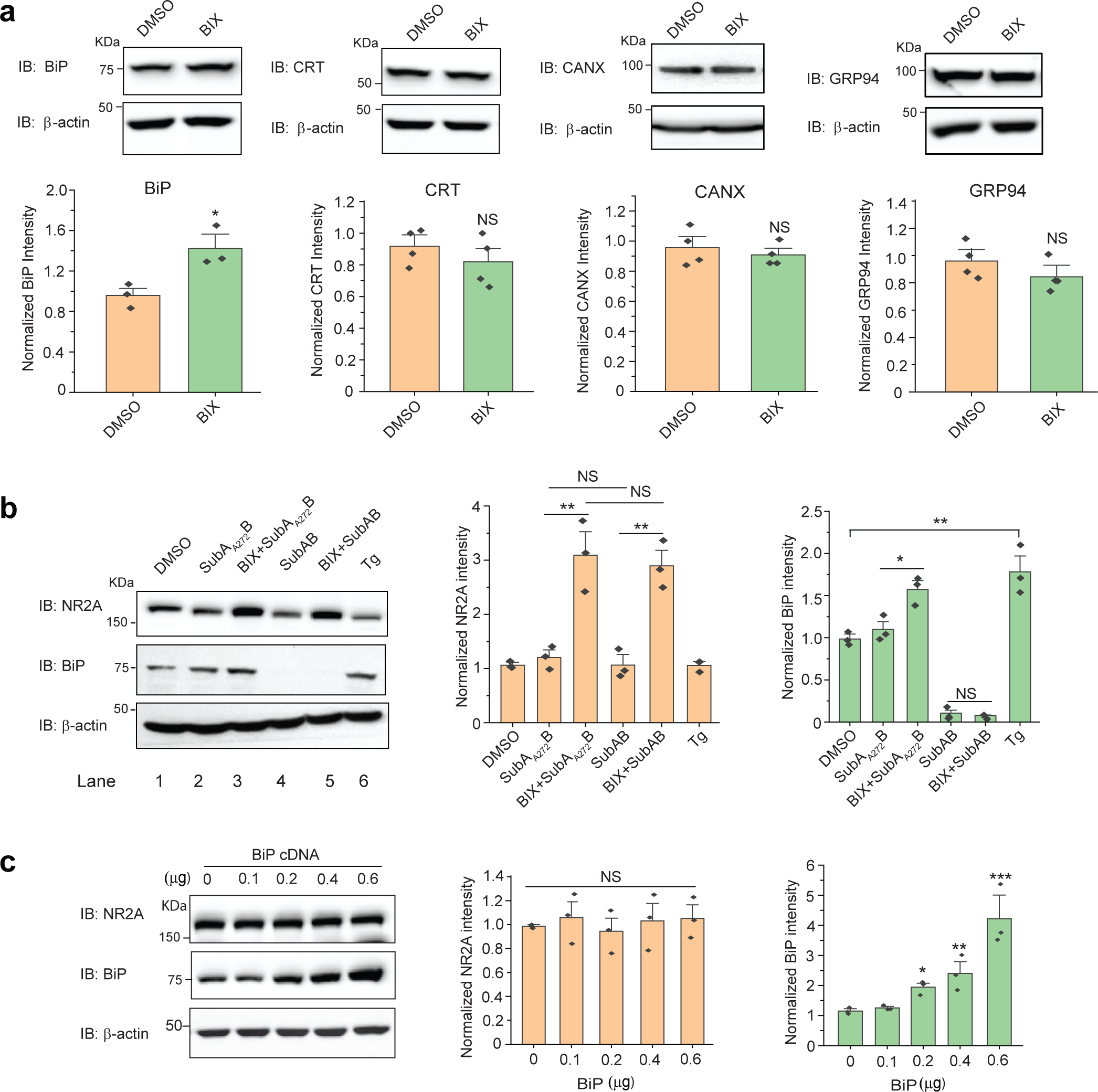
**Effect of BIX on increasing the NR2A variant total protein level is not dependent on BiP. a**, Effect of BIX (10 μM, 24 h) on the ER proteostasis network components, including BiP, CRT, CANX, and GRP94, in HEK293T cells stably expressing NR1NR2A(M705V). CRT: calreticulin. CANX: calnexin. Quantifications of the normalized band intensities were shown on the lower panels. **b**, Effect of BiP inhibition on BIX’s effect to increase the total expression levels of NR2A and BiP in HEK293T cells stably expressing NR1NR2A(M705V) according to western blot analysis. SubAB (0.5 μg/mL, 6 h), a potent BiP-specific protease, was applied to cell culture media to deplete BiP. SubAA272B served as a negative control of SubAB. Tg (thapsigargin) served as a positive control to increase the BiP protein level. **c**, Effect of BiP overexpression on the total protein levels of NR2A in HEK293T cells stably expressing NR1NR2A(M705V). Each data point is reported as mean ± SEM. Student’s t-test (for comparison of two groups) or one-way ANOVA followed by a post-hoc Tukey test (for comparison of three or more groups) was used for statistical analysis. *, p < 0.05; **, p < 0.01; ***, p < 0.001.

We next determined whether BIX’s effect on increasing NR2A(M705V) protein level is dependent on BiP. SubAB is a bacterial toxin with protease activity that selectively cleaves BiP, and SubAA272B is a negative control SubAB mutant that lacks BiP-specific protease activity [41]. Western blot analysis confirmed that BiP was dramatically depleted with SubAB treatment (0.5 μg/mL, 6 h); however, such BiP depletion did not affect the NR2A(M705V) protein levels (**Fig. 4b**. cf. lane 4 to 2). In addition, when BiP was depleted with SubAB, BIX treatment still significantly increased NR2A(M705V) protein levels (**Fig. 4b**. cf. lane 5 to 4). Furthermore, we carried out the BiP depletion with SubAB post BIX treatment (10 µM, 24 h) and demonstrated that BiP depletion did not diminish the increased NR2A(M705V) protein levels afforded by BIX treatment (**Fig. 4b**. cf. lane 5 to 3). In addition, we observed that BiP overexpression (up to 4.2- fold) did not change the NR2A(M705V) subunit protein levels (**Fig. 4c**). Therefore, these results indicated that although BIX upregulated BiP protein levels, the BIX-mediated increase of NR2A(M705V) protein levels did not principally rely on the BIX-upregulated BiP protein level. Other cellular signaling pathways must be involved in the regulation of the activity of BIX on NR2A(M705V) subunits.

Since the UPR is a major cellular pathway to enhance the ER folding capacity to handle misfolded proteins, we sought to determine whether BIX utilized the UPR to correct the folding and trafficking of NR2A(M705V). The UPR contains three signaling arms, namely IRE1, ATF6, and PERK, which are all ER integral membrane proteins. Activation of IRE1 and ATF6 is mainly cytoprotective by upregulating the expression of molecular chaperones to handle misfolded proteins, whereas activation of PERK often leads to apoptosis [24]. Upon activation, IRE1 oligomerizes, leading to the autophosphorylation of its kinase domain and the splicing of the mRNA of XBP1. Spliced XBP1 (XBP1s) is a potent transcription factor that promotes the ER folding capacity. Western blot analysis demonstrated that BIX treatment significantly increased the protein level of XBP1s (**Fig. 5a**), indicating the BIX modestly activated the IRE1 pathway. Upon activation, ATF6, a type-II membrane protein, is translocated form the ER to the Golgi and processed by S1P and S2P proteases to release the N-terminal cytoplasmic fragment of ATF6 (ATF6-N), which acts as a transcription factor to enhance ER folding. Western blot analysis showed that BIX treatment increased the protein level of the cleaved ATF6-N (**Fig. 5a**), as an indication of the activation of the ATF6 pathway. Upon activation, PERK phosphorylates eIF2α (eukaryotic translation initiation factor 2α) to reduce global translation, but eIF2α phosphorylation selectively induces the expression of ATF4 and its downstream target CHOP, a proapoptotic transcription factor. Western blot analysis showed that BIX treatment did not change the protein level of CHOP (**Fig. 5a**), indicating that the PERK arm was not activated. Thapsigargin (Tg), a well-established UPR activator, activated all three arms of the UPR, but did not increase the NR2A(M705V) protein level in HEK293T cells (**Fig. 5a**).

**Figure 5.**
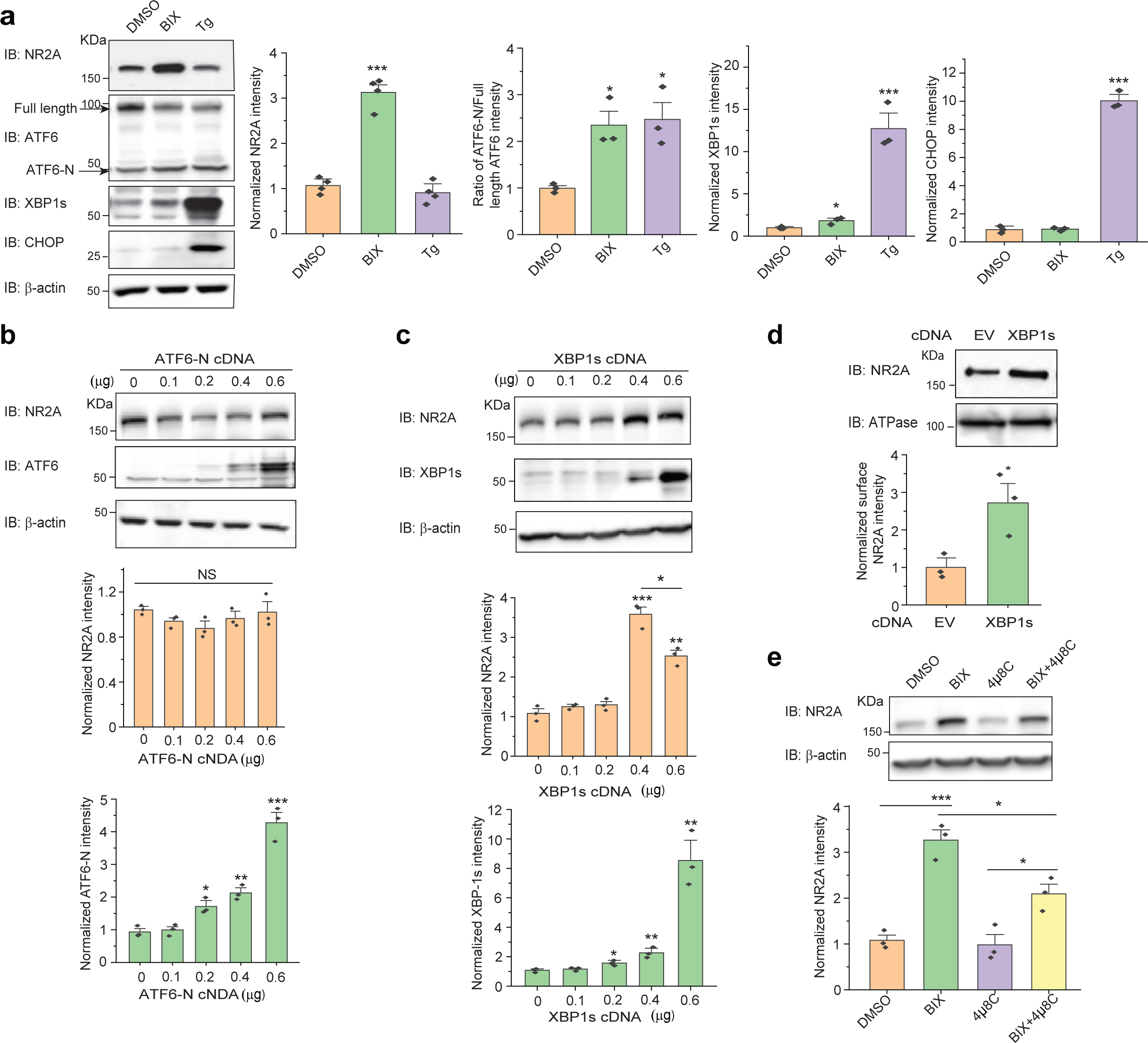
**BIX increases variant NR2A subunits expression through pharmacological activation of the unfolded protein response (UPR). a**, Effect of BIX (10 μM, 24 h) on the total protein levels of NR2A and UPR-associated proteins, including ATF6-N, XBP-1s and CHOP, in HEK293T cells stably expressing NR1NR2A(M705V). Tg (thapsigargin) serves as a positive control to activate the UPR. **b**, and **c**, Effect of ATF6-N (**b**) and XBP-1s (**c**) overexpression on the total protein levels of NR2A in HEK293T cells stably expressing NR1NR2A(M705V). **d**, XPB-1s overexpression enhances surface protein levels of NR2A in HEK293T cells stably expressing NR1NR2A(M705V). Na^+^/K^+^-ATPase serves as the loading control of surface protein. **e,** Effect of an IRE1 inhibitor 4µ8C (32 µM, 24h) on the BIX (10 µM, 24h)-induced protein levels of NR2A in HEK293T cells stably expressing NR1NR2A(M705V). Each data point is reported as mean ± SEM. Student’s t-test (for comparison of two groups) or one-way ANOVA followed by a post-hoc Tukey test (for comparison of three or more groups) was used for statistical analysis. *, p < 0.05; **, p < 0.01; ***, p < 0.001.

Since BIX treatment modestly activated the IRE1 and ATF6 arms of the UPR, we continued to determine the effect of genetic activation of ATF6 and IRE1 on the maturation of the NR2A(M705V) subunits. Overexpression of active ATF6-N up to 4.2-fold of the endogenous level did not increase the total protein expression levels of NR2A(M705V) (**Fig. 5b**), suggesting that ATF6 activation did not contribute to the effect of BIX on increasing NR2A(M705V) protein level. In contrast, western blot analysis showed that the genetic activation of the IRE1 pathway by overexpressing XBP1s resulted in a substantial increase of the total protein level of NR2A(M705V) subunits (**Fig. 5c**, cf. lanes 4 and 5 to lane 1). Moreover, it appeared that the modest overexpression of XBP1s at 2.1-fold to endogenous level, which is similar to the level that BIX induced, resulted in the most dramatic increase of NR2A(M705V) protein (**Fig. 5c**, cf. lane 4 to 5). In addition, surface biotinylation experiments showed that modest XBP-1s overexpression substantially increased the surface protein level of the NR2A(M705V) subunits (**Fig. 5d**). To further determine whether the BIX’s effect depended on the activation of the IRE1 pathway, we co-applied 4μ8C, a specific IRE1 inhibitor acting on the RNase domain [42] with BIX treatment. Western blot analysis demonstrated that inhibiting the IRE1 pathway significantly reduced the NR2A(M705V) protein levels that was induced by BIX treatment (**Fig. 5e**, cf. lane 4 to 2), indicating that indeed the IRE1 activation contributed to the effect of BIX on enhancing NR2A(M705V) proteostasis. The differentiating effects of ATF6 and IRE1 on the NR2A(M705V) variant protein levels could be due to the distinct downstream gene targets between ATF6 activation and IRE1 activation, which would adapt the ER proteostasis differently [43].

In summary, the above experiments clearly demonstrated that modest IRE1/XBP1s activation enhanced the surface trafficking of NR2A(M705V). Therefore, through activating the IRE1/XBP1s pathway, BIX enhances the folding, trafficking, and function of the NR2A(M705V) variant.

## Discussion

Our data corroborate that BIX enhances surface trafficking of epilepsy-associated NR2A variant subunits of NMDARs, including NR2A(M705V) and NR2A(A727T). Once they reach the plasma membrane, the function of the variant receptors is restored (**Fig. 2f**). Our results support the mechanism for BIX-mediated rescue of misfolding-prone NR2A variant NMDARs proposed in **Fig. 6**. BIX treatment modestly activates the IRE1/XBP1s arm of the UPR pathway to remodel the ER proteostasis network, including the regulation of molecular chaperones [43], which leads to enhanced ER folding capacity. This operation further stabilizes the misfolding-prone NR2A variant, inhibits its ERAD, enhances its folding and assembly in the ER membrane, and promotes the productive trafficking of the variant-containing NMDARs to the Golgi and further onward to the plasma membrane. Consequently, BIX treatment corrects the function of the variant-containing NMDARs by enhancing their trafficking to the plasma membrane.

**Figure 6.**
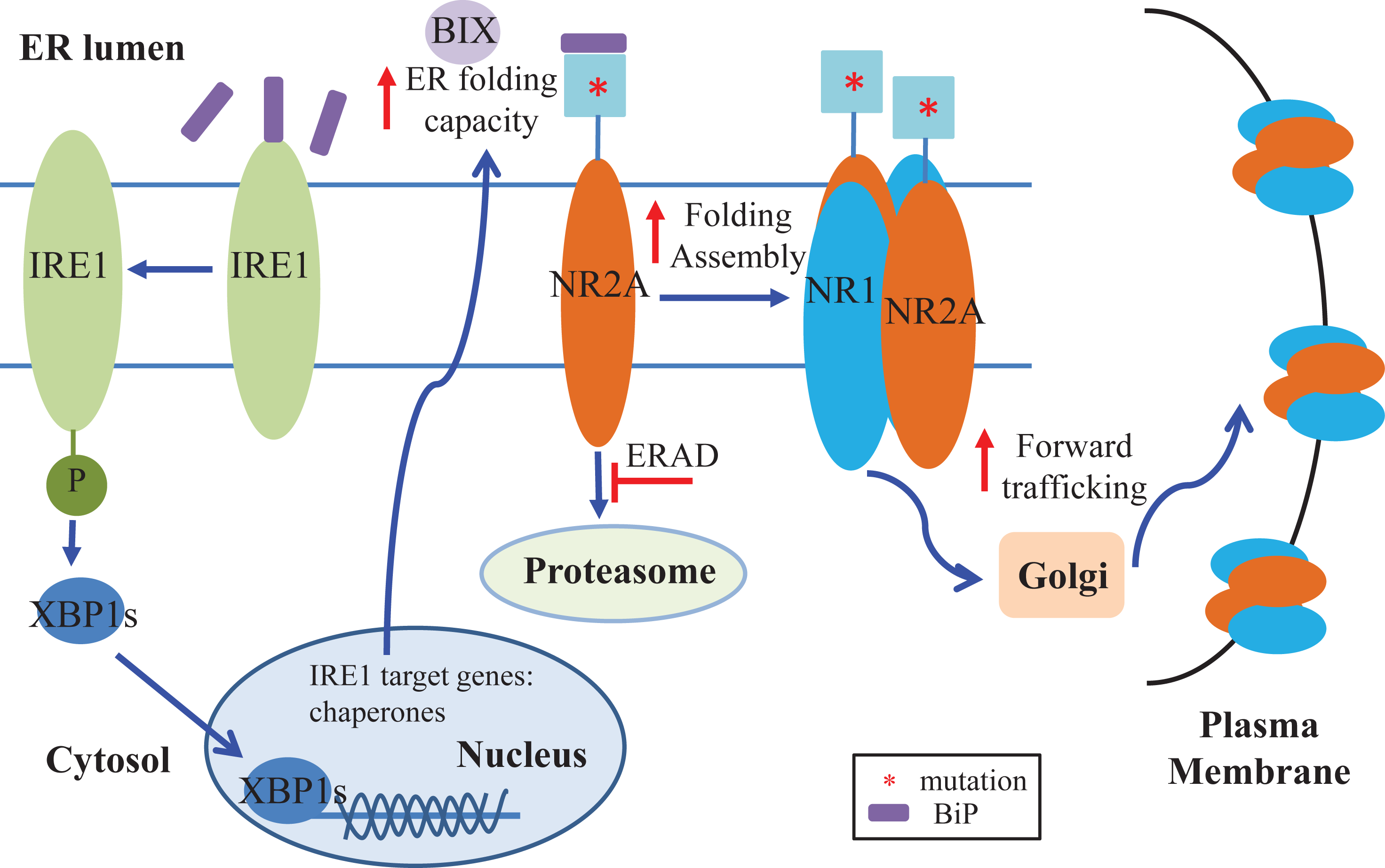
Mechanism of the effect of BIX on NMDAR proteostasis. Our working model presents BIX’s capability to enhance the proteostasis network of NMDARs harboring NR2A mutations by activation of the IRE1 pathway to enhance ER folding capacity and to attenuate variant NR2A degradation through ERAD. Activation of the IRE1 arm of the UPR results in the splicing of XBP1, which in turn acts as a transcription factor to upregulate genes associated with protein folding, assembly, trafficking, and degradation [53]. Together, enhanced folding and assembly and reduction of ERAD of variant NR2A subunits result in an increased population of mature NMDARs that can traffic through the Golgi and to the plasma membrane to perform their function as ion channels. Asterisk (*) represents mutations on the NR2A subunits.

Increasing reports have investigated the effects of missense mutations on NR2A protein function in various expression systems [44–46], a common outcome of which is that epilepsy-associated NR2A variants have strikingly variable functional consequences on NMDARs [10]. The significance of NMDARs is well-demonstrated by the range of clinical phenotypes of patients with variants in the genes encoding NR2A subunits. A standard approach for rescuing loss-of-function phenotypes is the usage of pharmacological modulators to enhance receptor function. The ability of ligand binding to control ion-channel gating has been shown to be a functional checkpoint for glutamate receptors, including NMDARs, to exit the ER [13, 14, 47]. Thereby, we used epilepsy-associated M705V and A727T variations, which are located in the LBD of the NR2A subunits of NMDARs, to evaluate how BIX remodels the ER proteostasis network to rescue the misfolding-prone variants of NMDARs.

We first determined the protein expression level and functional characteristics of NMDARs containing M705V or A727T variants in the NR2A subunits. The results from transfected HEK293T cells revealed that total and surface protein levels of NR2A were significantly reduced in both M705V and A727T variants (**Fig. 1c-1e**). The results coincided with the reduced peak currents recorded by whole-cell patch-clamp technique for NR1NR2A(M705V) and NR1NR2A(A727T) (**Fig. 1f**). Moreover, the variants in NR2A subunits showed the increased degradation by the proteasome pathway with the application of MG132 compared to NR2A WT (**Fig. 1g**). The results ascertain that NMDARs harboring M705V or A727T variants in the NR2A subunits are prone to misfolding in the ER and inefficient trafficking to the cell membrane.

In the past, researchers attempted to validate BiP as a potentially potent therapeutic target for pharmacological manipulation of neuronal cell damage in the brain[48]. Previous reports revealed that treatment with BIX had anti-apoptotic and neuroprotective effects against neuronal cell damage in animal models [49–51], which implied that BIX may be developed as a promising candidate to attenuate ER stress for the treatment of ER stress-associated diseases. Our results showed that BIX treatment not only increased the total protein levels but also surface protein levels of NR2A subunits containing M705V or A727T variant (**Fig. 2a-2e**), indicating that BIX promotes the folding, forward trafficking, and importantly functional surface expression of NR1NR2A(M705V) and NR1NR2A(A727T). The time course study of single-dose BIX treatment showed the optimal rescue level was achieved at 12-24h (**Fig. 2c**), which may be possibly due to the involvement of protein maturation, such as protein folding and trafficking. Furthermore, the increased currents induced by BIX treatment in NMDARs containing NR2A variants were comparable to that recorded in NR1NR2A WT NMDARs, which suggested that BIX could potentially improve the functional rescue of malignant NR2A variants (**Fig. 2f**). These results provided evidence that the misfolding and trafficking deficiency of NMDARs harboring NR2A variants can be restored by BIX treatment.

We sought to investigate how BIX adapts the ER proteostasis network to correct NR2A variant function. Among the major ER chaperones tested, BIX treatment did not significantly change the protein levels of calnexin, calreticulin, and GRP94, but increased BiP levels in HEK293T cells stably expressing NR1NR2A(M705V) (**Fig. 4a**). Interestingly, such BIX- upregulated BiP expression did not seem to affect BIX’s effect on the variant NR2A subunits (**Fig. 4b-4c**). Since the UPR is a major pathway to regulate ER proteostasis, we next determined the effect of BIX treatment on the activation of the UPR. BIX activated both the IRE1 and ATF6 arms of the UPR. However, IRE1 activation by XBP-1s overexpression significantly increased the total protein level of NR2A(M705V) subunits in HEK293T cells, whereas ATF6 overexpression did not (**Fig. 5b-5c**). The differing effects between IRE1 and ATF6 activation on the NR2A variants is likely due to the fact that they have different downstream targets. These results suggest that the activation of the IRE1 pathway rather than the ATF6 pathway plays an important role in the BIX-induced rescue of misfolding-prone NR2A variant receptors. It merits further investigation to find out which downstream targets of the IRE1 activation play a key role in correct the misfolding of NR2A variants.

In conclusion, we demonstrated that the BIX treatment enhances the productive folding and forward trafficking, and further restores the function of NMDARs containing pathogenic variant NR2A subunits by modestly activating the IRE1 pathway. This study provides a proof-of-concept case about remodeling ER proteostasis to rescue the loss-of-function NR2A variant NMDARs associated with protein misfolding diseases. Such a folding-correction strategy has a dual role in the functional rescue of NMDARs: first, it will increase the number of the functional channels on the plasma membrane to enhance the post-synaptic activities; second, the increased surface receptors will be available to current known NMDAR-specific ligands, such as agonists and positive allosteric modulators, providing a potential synergistic action between two mechanistically distinct compounds.

## Methods

### Plasmids, chemicals, and antibodies

The pcDNA3.1-NR2A (catalog #: OHu24642D, NM_000833, human) and the pcDNA3.1-NR1 (catalog #: OHu22255D, NM_007327, human) plasmids were purchased from GenScript. The pcDNA3.1-BiP plasmid was provided by Dr. Tohru Mizushima (Kumamoto University). N-terminal HA-tagged full-length pCGN-ATF6-N plasmid came from Addgene (catalog #: 11974, human). The XBP1s plasmid was a kind gift from Dr. Richard N. Sifers (Baylor College of Medicine).

BIX (BiP protein inducer X) was obtained from Sigma (catalog #: SML1073). Thapsigargin (Tg) was obtained from Enzo Life Sciences (catalog #: BML-PE180-0001). (+)- MK801 maleate was purchased from Hellobio (catalog #: HB0004). Resazurin was obtained from MP Biomedicals (catalog #: 0219548101). SubAB, a bacterial toxin with protease activity highly selective for BiP and SubAA272B, a non-proteolytic negative control for SubAB, were prepared according to the published procedure [41]. 4µ8C (8-formyl-7-hydroxy-4- methylcoumarin), a specific IRE1 inhibitor, was purchased from Cayman Chemical (catalog #: 22110). Other chemicals were purchased from Sigma unless stated otherwise.

The rabbit polyclonal anti-NR2A (catalog #: ab124913; 1:3,000), mouse polyclonal anti- NR2A (catalog #: ab240884; 1:100), and rabbit monoclonal anti-Na^+^/K^+^-ATPase antibody (catalog #: ab76020; 1:3,000) were purchased from Abcam. The mouse polyclonal anti-calreticulin (catalog #: ADI-SPA-601; 1:3,000), the rabbit polyclonal anti-calnexin (CANX) (catalog #: ADI-SPA-860-F, 1:5,000), and the rat monoclonal anti-GRP94 (clone 9G10) (catalog #: ADI-SPA-850-F, 1:1,000) antibodies were purchased from Enzo Life Sciences. The rabbit polyclonal anti-XBP1 antibody (M-186) is from Santa Cruz (catalog #: SC-7160, 1:1,000). The mouse monoclonal anti-β-actin antibody (catalog #: A1978, 1:10,000) was obtained from Sigma Aldrich. The rabbit polyclonal anti-BiP antibody is from Abgent (catalog #: AP50016, 1:5,000). The rabbit monoclonal anti-ATF6 antibody (catalog #: 24169-1-AP; 1:3,000) was from Proteintech. The mouse polyclonal anti-CHOP antibody was obtained from Cell Signaling (catalog #: 2895).

The NR2A(M705V) or NR2A(A727T) mutation was introduced to the NR2A subunit using QuikChange II site-directed mutagenesis Kit (Agilent Genomics). DNA sequencing was used to confirm the cDNA sequences. The forward and reverse primers for NR2A(M705V) are 5’-CCTTTCTGATTAAATTTGGTCACGTACTGATGCATGTAGGGATAG-3’ and 5’-CTATC CCTACATGCATCAGTACGTGACCAAATTTAATCAGAAAGG-3’; the forward and reverse primers for NR2A(A727T) are 5’-GCATCGTAGATGAAAGTGTCCAGCTTCCCCGTT-3’ and 5’-AACGGGGAAGCTGGACACTTTCATCTACGATGC-3’.

### Cell culture and transfection

HEK293T cells (catalog #: CRL-3216) were obtained from ATCC. Human A2780 cells (catalog #: 93112519) were purchased from Sigma. Cells were cultured in Dulbecco’s Modified Eagle Medium (DMEM) (Fisher, catalog #: SH3024301) containing 10% heat-inactivated fetal bovine serum (ThermoFisher, catalog #: SH3039603HI) and 1% Penicillin-Streptomycin (Hyclone, catalog #: sv30010) at 37°C in 5% CO2. Cells were seeded in 10-cm dishes or 6-well plates and allowed to reach ∼70% confluency before transient transfection using TransIT-2020 (Mirus, catalog #: MIR 5400) according to the manufacturer’s instruction. 50 µM MK801 (hellobio catalog #: HB0004) and 2.5 mM MgCl2 were added to medium 4h post transient transfection to avoid excitatory toxicity. Then, cells were harvested for further analysis after transfection for 24h.

Stable cell lines for NR1NR2A, NR1NR2A(M705V) and NR1NR2A(A727T) were generated using the G418 selection method in the presence of MK801 and MgCl2 to block the receptors as they were expressed. Briefly, cells were transfected with NR1: NR2A (1:1), NR1: NR2A(M705V) (1:1) or NR1: NR2A(A727T) (1:1) plasmids, selected in DMEM supplemented with 1.0 mg/mL G418 (Enzo Life Sciences, catalog#: ALX-380-013) for 10 days, and then maintained in DMEM supplemented with 0.4 mg/mL G418. G418 resistant cells expressing NMDARs were verified by western blot analysis and then used for experiments.

### SDS-PAGE and western blot analysis

Cells were harvested with cold Dulbecco’s phosphate-buffered saline (DPBS) (Fisher, catalog#: SH3002803) into a centrifuge tube and then lysed with lysis buffer (50mM Tris, pH 7.5, 150mM NaCl, and 2 mM Dodecyl-B-D-maltoside (DDM) (GoldBio, catalog #: DDM5) supplemented with complete protease inhibitor cocktail (Roche #04693116001). Cells were vortexed and ultrasonicated for 40 s three times. Lysates were cleared at 21,000 ×g, 4 °C for 10 min. Protein concentrations were measured by Thermo Fisher MicroBCA kit (ThermoFisher Pierce #23235). Cell lysates were loaded with 4 × Laemmli buffer (Biorad, catalog #: 1610747) including 2-Mercaptoethanol (1:10 v/v; Sigma #M3148) and subjected to 4-20% SDS-PAGE gel. Protein ladder (Biorad, catalog #: 1610395) was used to locate the molecular weight. Western blot analysis was carried out using appropriate antibodies with dilutions as listed above. Band intensity was quantified using ImageJ software. The β-actin and the Na^+^/K^+^-ATPase serve individually as a loading control of total protein and plasma membrane proteins respectively. The total protein was first normalized to the loading control and then the vehicle, DMSO, or WT control.

### Biotinylation of cell surface proteins

HEK293T cells stably expressing NMDAR variants were plated in Poly-L-Lysine pre-coated 10-cm dishes for surface biotinylation experiments according to published procedure [54]. In brief, cells were transfected with the corresponding NR1NR2A plasmids for 24 h, and then treated with BIX at 10 μM for another 24h. The medium was supplemented with antagonists to decrease the cell excitatory toxicity. Intact cells were rinsed gently twice with ice-cold DPBS and incubated with the membrane-impermeable biotinylation reagent Sulfo-NHS SS-Biotin (0.5 mg/mL; ThermoFisher Pierce) in DPBS containing 1 mM CaCl2 and 0.5 mM MgCl2 (DPBS-CM) for 30 min on ice to label surface membrane proteins. 10 mM glycine in ice-cold PBS-CM was added to plates for 5 min at 4°C to quench the reaction. To block the sulfhydryl groups, 5 nM N-ethylmaleimide (NEM) in DPBS was added for 15 min at room temperature. Cells were scraped, transferred to a tube, and were solubilized at 4 °C overnight in homogenization buffer (Triton X- 100, 1%; Tris–HCl, 50 mM; NaCl, 150 mM; and EDTA, 5 mM; pH 7.5) supplemented with Roche protease inhibitor cocktail and 5 mM NEM. The supernatant containing the biotinylated surface proteins was obtained followed by centrifuging (16,000 ×g, 10 min at 4°C) the lysates. The concentration of the supernatant was measured using microBCA assay (ThermoFisher Pierce). Biotinylated surface proteins were affinity-purified by incubating the obtained supernatant for 1 h at 4°C with 40 μL immobilized neutravidin-conjugated agarose bead slurry (ThermoFisher Pierce, catalog #: 29201). The beads were kept and washed with wash buffer (Triton X-100, 0.5%; Tris–HCl, 50 mM; NaCl, 150 mM; and EDTA, 5 mM; pH 7.5) and 1X TBS, three times each. Surface proteins were eluted from beads with 70 μL elution buffer (2x Laemmli sample buffer with 100 mM DTT and 6 M urea; pH 6.8) by vortex 30 min at room temperature before being subjected to SDS-PAGE and western blot analysis.

### Confocal immunofluorescence

The labeling of surface NMDARs and confocal immunofluorescence microscopy analysis were performed according to published procedure [55]. Briefly, cells cultured on coverslips were fixed with 4% formaldehyde in DPBS for 15 min at 4°C, blocked with 10% goat serum for 30 min at room temperature, and incubated in 50 μL of 2% goat serum in PBS containing mouse monoclonal anti-NR2A antibody (abcam #240884, 1:200) and rabbit monoclonal anti- Na^+^/K^+^-ATPase, a plasma membrane marker (Abcam #ab76020, 1:200) for overnight at 4°C. The primary antibodies were used at 1:200 dilutions. The cells were incubated at room temperature with 50 μL an Alexa 488-conjugated goat anti-mouse secondary antibody (1:500, ThermoFisher, catalog #: A11001) and an Alexa 568-conjugated goat anti-rabbit secondary antibody (1:500, ThermoFisher, catalog #: A11037) for 1 h. Then, cells were rinsed with DPBS two times and incubated with DAPI (1 μg/mL) for 5 min to stain the nucleus. The coverslips were then mounted and sealed. For confocal immunofluorescence microscopy, an Olympus IX-81 Fluoview FV3000 confocal laser scanning system was used. A 40X objective was used to collect images using FV31S-SW software. Quantification of the fluorescence intensity was achieved using the ImageJ software.

### Cycloheximide-chase (CHX) assay

CHX assay was used to assess protein degradation and performed as previously mentioned [54]. Cycloheximide (100 µg/mL, Enzo life Science #ALX-380-269) was added to the culture medium to stop mRNA translation. Briefly, HEK293T cells expressing NR1NR2A (M705V) NMDARs were treated with DMSO or BIX for 24h, and then chased for the respective time points and harvested. Cells were lysed for total protein and then subjected to SDS-PAGE.

### Endoglycosidase H (endo H) enzyme digestion assay

Endo H assay was used to monitor the efficiency of protein trafficking from ER to Golgi. To remove asparaginyl-N-acetyl-D-glucosamine in the N-linked glycans incorporated on the NR2A subunit in the ER, the Endo H enzyme (NEBiolab, catalog #: P0703L) was used to digest the cell lysates at 37°C for 24h. The samples treated by the Peptide-N-Glycosidase F (PNGase F) enzyme (NEBiolab, catalog #: P0704L) serve as a control for unglycosylated NR2A subunits. Treated samples were then subjected to SDS-PAGE and western blot analysis.

### Automated patch-clamping recording with IonFlux Mercury 16 instrument

Whole-cell currents of NMDARs composed of NR1NR2A were recorded 48 h post-transfection of HEK293T cells. Automated patch-clamping was performed on the Ionflux Mercury 16 instrument (Fluxion Biosciences, California). The extracellular solution (ECS) contained the following: 145 mM NaCl, 4 mM KCl, 2 mM CaCl2(2H2O), 10 mM glucose, 10 mM HEPES (pH adjusted with NaOH). The intracellular solution (ICS) contained the following: 135 mM CsCl, 1 mM EGTA, 5 mM CsOH, 10 mM HEPES, 10 mM NaCl (pH adjusted with CsOH). In brief, cells were grown to ∼70% confluence on 10-cm dishes and rinsed with DPBS. Then 3 mL Accutase (Sigma Aldrich, catalog #: A6964) was added to the cells and incubated for 2 min at 37 °C until the cells were floating with minimal clumps under a microscope. Cells were pelleted via centrifugation for 1 min at 200 ×g, supernatant was aspriated and cells were resuspended in serum-free medium HEK293T SFM II (Gibco, catalog #: 11686-029), supplemented with 25 mM HEPES (Gibco, catalog #: 15630-080) and 1% penicillin streptomycin (Hyclone, catalog #: sv30010). Cells were gently shaken at room temperature for at least 30 min before experiments. Mercury 16 plates were prepared according to manufacture’s procedure. Whole-cell NMDARs currents were recorded at a holding potential of -60 mV at the application of 10 mM glutamate and 100 µM glycine as indicated. The data were acquired and analyzed by Fluxion Data Analyzer.

### Resazurin cell toxicity assay

Resazurin cell toxicity assay was performed colorimetrically as previously described [37]. HEK293T cells stably expressing NR1NR2A (M705V) NMDARs were seeded into a 96-well microtitre plate. The cells were separated into nine groups which were treated with DMSO or BIX (0.5, 1, 2.5, 5, 10, 20 or 40 μM) for 24 h, or thapsigargin (2 μM, 6 h). Resazurin stain (10 μL of 0.15 mg/mL in DPBS) was added to each well using a multichannel auto pipette and re-incubated the plate at 37 °C for 2 h before plate reading. The fluorescence signal to proportion cell metabolism/viability was determined spectrophotometrically at excitation 530 nm using a reference wavelength of emission 590 nm with a microplate reader (Tecan - Spark, A-5082, Grödig, Austria).

### Quantification and statistical analysis

All values were presented as mean ± SEM. Data for statistical analysis was evaluated by using a two-tailed Student’s t-test between two groups as appropriate, and the analysis of variance (ANOVA) followed by a post-hoc Tukey test was applied for comparisons in multiple groups. A p < 0.05 was defined as statistically significant. ∗, p< 0.05; ∗∗, p< 0.01; ∗∗∗, p< 0.001.

## SUPPLEMENTAL INFORMATION

Supplemental Information includes two figures.

## ACKNOWLEDGMENTS

This work was supported by the National Institutes of Health (R01NS117176 to TM).

## AUTHOR CONTRIBUTIONS

Conceptualization, PZ, TM, and YW; Data curation: PZ, and YW; Formal analysis: PZ, TM, and YW; Funding acquisition: TM; Supervision: TM and YW; Writing – original draft: PZ and YW; Writing – review & editing: PZ, TB, JP, AP, TM, and YW.

## References

1 Hansen KB, Wollmuth LP, Bowie D, Furukawa H, Menniti FS, Sobolevsky AI, et al. Structure, Function, and Pharmacology of Glutamate Receptor Ion Channels. Pharmacological reviews 2021; 73: 298–487.

2 Yong XLH, Zhang L, Yang L, Chen X, Tan JZA, Yu X, et al. Regulation of NMDA receptor trafficking and gating by activity-dependent CaMKIIalpha phosphorylation of the GluN2A subunit. Cell reports 2021; 36: 109338.

3 Karakas E, Furukawa H. Crystal structure of a heterotetrameric NMDA receptor ion channel. Science 2014; 344: 992–7.

4 Lee CH, Lü W, Michel JC, Goehring A, Du J, Song X, et al. NMDA receptor structures reveal subunit arrangement and pore architecture. Nature 2014; 511: 191–7.

5 Sanz-Clemente A, Nicoll RA, Roche KW. Diversity in NMDA receptor composition: many regulators, many consequences. The Neuroscientist : a review journal bringing neurobiology, neurology and psychiatry 2013; 19: 62–75.

6 Vieira M, Yong XLH, Roche KW, Anggono V. Regulation of NMDA glutamate receptor functions by the GluN2 subunits. Journal of neurochemistry 2020; 154: 121–43.

7 Stroebel D, Casado M, Paoletti P. Triheteromeric NMDA receptors: from structure to synaptic physiology. Current opinion in physiology 2018; 2: 1–12.

8 XiangWei W, Jiang Y, Yuan H. De Novo Mutations and Rare Variants Occurring in NMDA Receptors. Current opinion in physiology 2018; 2: 27–35.

9 Myers SJ, Yuan H, Kang JQ, Tan FCK, Traynelis SF, Low CM. Distinct roles of GRIN2A and GRIN2B variants in neurological conditions. F1000Research 2019; 8(F1000 Faculty Rev): 1940.

10 Elmasri M, Hunter DW, Winchester G, Bates EE, Aziz W, Van Der Does DM, et al. Common synaptic phenotypes arising from diverse mutations in the human NMDA receptor subunit GluN2A. Communications biology 2022; 5: 174.

11 Benske TM, Mu TW, Wang YJ. Protein quality control of N-methyl-D-aspartate receptors. Frontiers in cellular neuroscience 2022; 16: 907560.

12 Fu YL, Wang YJ, Mu TW. Proteostasis Maintenance of Cys-Loop Receptors. Advances in protein chemistry and structural biology 2016; 103: 1–23.

13 Swanger SA, Chen W, Wells G, Burger PB, Tankovic A, Bhattacharya S, et al. Mechanistic Insight into NMDA Receptor Dysregulation by Rare Variants in the GluN2A and GluN2B Agonist Binding Domains. American journal of human genetics 2016; 99: 1261–80.

14 She K, Ferreira JS, Carvalho AL, Craig AM. Glutamate binding to the GluN2B subunit controls surface trafficking of N-methyl-D-aspartate (NMDA) receptors. The Journal of biological chemistry 2012; 287: 27432–45.

15 Mah SJ, Cornell E, Mitchell NA, Fleck MW. Glutamate receptor trafficking: endoplasmic reticulum quality control involves ligand binding and receptor function. The Journal of neuroscience : the official journal of the Society for Neuroscience 2005; 25: 2215–25.

16 Addis L, Virdee JK, Vidler LR, Collier DA, Pal DK, Ursu D. Epilepsy-associated GRIN2A mutations reduce NMDA receptor trafficking and agonist potency - molecular profiling and functional rescue. Scientific reports 2017; 7: 66.

17 Needham PG, Guerriero CJ, Brodsky JL. Chaperoning Endoplasmic Reticulum-Associated Degradation (ERAD) and Protein Conformational Diseases. Cold Spring Harbor perspectives in biology 2019; 11.

18 Adams BM, Oster ME, Hebert DN. Protein Quality Control in the Endoplasmic Reticulum. The protein journal 2019; 38: 317–29.

19 Sun Z, Brodsky JL. Protein quality control in the secretory pathway. The Journal of cell biology 2019; 218: 3171–87.

20 Grandjean JMD, Wiseman RL. Small molecule strategies to harness the unfolded protein response: where do we go from here? The Journal of biological chemistry 2020; 295: 15692–711.

21 Kelly JW. Pharmacologic Approaches for Adapting Proteostasis in the Secretory Pathway to Ameliorate Protein Conformational Diseases. Cold Spring Harbor perspectives in biology 2020; 12: a034108.

22 Marciniak SJ, Chambers JE, Ron D. Pharmacological targeting of endoplasmic reticulum stress in disease. Nature reviews. Drug discovery 2022; 21: 115–40.

23 Otero JH, Lizák B, Hendershot LM. Life and death of a BiP substrate. Seminars in cell & developmental biology 2010; 21: 472–8.

24 Walter P, Ron D. The unfolded protein response: from stress pathway to homeostatic regulation. Science 2011; 334: 1081–6.

25 Hetz C, Zhang K, Kaufman RJ. Mechanisms, regulation and functions of the unfolded protein response. Nature reviews. Molecular cell biology 2020; 21: 421–38.

26 Wiseman RL, Mesgarzadeh JS, Hendershot LM. Reshaping endoplasmic reticulum quality control through the unfolded protein response. Molecular cell 2022; 82: 1477–91.

27 Gorbatyuk MS, Gorbatyuk OS. The Molecular Chaperone GRP78/BiP as a Therapeutic Target for Neurodegenerative Disorders: A Mini Review. Journal of genetic syndromes & gene therapy 2013; 4: 128.

28 Kudo T, Kanemoto S, Hara H, Morimoto N, Morihara T, Kimura R, et al. A molecular chaperone inducer protects neurons from ER stress. Cell death and differentiation 2008; 15: 364–75.

29 Huntley GW, Vickers JC, Morrison JH. Cellular and synaptic localization of NMDA and non-NMDA receptor subunits in neocortex: organizational features related to cortical circuitry, function and disease. Trends in neurosciences 1994; 17: 536–43.

30 Meador-Woodruff JH, Healy DJ. Glutamate receptor expression in schizophrenic brain. Brain research. Brain research reviews 2000; 31: 288–94.

31 Bekkers JM, Stevens CF. NMDA and non-NMDA receptors are co-localized at individual excitatory synapses in cultured rat hippocampus. Nature 1989; 341: 230–3.

32 Wong EH, Kemp JA, Priestley T, Knight AR, Woodruff GN, Iversen LL. The anticonvulsant MK-801 is a potent N-methyl-D-aspartate antagonist. Proceedings of the National Academy of Sciences of the United States of America 1986; 83: 7104–8.

33 Lemke JR, Lal D, Reinthaler EM, Steiner I, Nothnagel M, Alber M, et al. Mutations in GRIN2A cause idiopathic focal epilepsy with rolandic spikes. Nature genetics 2013; 45: 1067–72.

34 Liu XR, Xu XX, Lin SM, Fan CY, Ye TT, Tang B, et al. GRIN2A Variants Associated With Idiopathic Generalized Epilepsies. Frontiers in molecular neuroscience 2021; 14: 720984.

35 Carpenter AE, Jones TR, Lamprecht MR, Clarke C, Kang IH, Friman O, et al. CellProfiler: image analysis software for identifying and quantifying cell phenotypes. Genome biology 2006; 7: R100.

36 Valley CC, Cembran A, Perlmutter JD, Lewis AK, Labello NP, Gao J, et al. The methionine-aromatic motif plays a unique role in stabilizing protein structure. The Journal of biological chemistry 2012; 287: 34979–91.

37 Fu YL, Han DY, Wang YJ, Di XJ, Yu HB, Mu TW. Remodeling the endoplasmic reticulum proteostasis network restores proteostasis of pathogenic GABAA receptors. PloS one 2018; 13: e0207948.

38 North WG, Liu F, Tian R, Abbasi H, Akerman B. NMDA receptors are expressed in human ovarian cancer tissues and human ovarian cancer cell lines. Clinical pharmacology : advances and applications 2015; 7: 111–7.

39 Kohli E, Causse S, Baverel V, Dubrez L, Borges-Bonan N, Demidov O, et al. Endoplasmic Reticulum Chaperones in Viral Infection: Therapeutic Perspectives. Microbiology and molecular biology reviews : MMBR 2021; 85: e0003521.

40 Ni M, Lee AS. ER chaperones in mammalian development and human diseases. FEBS letters 2007; 581: 3641–51.

41 Paton AW, Beddoe T, Thorpe CM, Whisstock JC, Wilce MC, Rossjohn J, et al. AB5 subtilase cytotoxin inactivates the endoplasmic reticulum chaperone BiP. Nature 2006; 443: 548–52.

42 Cross BC, Bond PJ, Sadowski PG, Jha BK, Zak J, Goodman JM, et al. The molecular basis for selective inhibition of unconventional mRNA splicing by an IRE1-binding small molecule. Proceedings of the National Academy of Sciences of the United States of America 2012; 109: E869–78.

43 Shoulders MD, Ryno LM, Genereux JC, Moresco JJ, Tu PG, Wu C, et al. Stress-independent activation of XBP1s and/or ATF6 reveals three functionally diverse ER proteostasis environments. Cell reports 2013; 3: 1279–92.

44 Yuan H, Hansen KB, Zhang J, Pierson TM, Markello TC, Fajardo KV, et al. Functional analysis of a de novo GRIN2A missense mutation associated with early-onset epileptic encephalopathy. Nature communications 2014; 5: 3251.

45 Sibarov DA, Bruneau N, Antonov SM, Szepetowski P, Burnashev N, Giniatullin R. Functional Properties of Human NMDA Receptors Associated with Epilepsy-Related Mutations of GluN2A Subunit. Frontiers in cellular neuroscience 2017; 11: 155.

46 Chen W, Tankovic A, Burger PB, Kusumoto H, Traynelis SF, Yuan H. Functional Evaluation of a De Novo GRIN2A Mutation Identified in a Patient with Profound Global Developmental Delay and Refractory Epilepsy. Molecular pharmacology 2017; 91: 317–30.

47 Gill MB, Vivithanaporn P, Swanson GT. Glutamate binding and conformational flexibility of ligand-binding domains are critical early determinants of efficient kainate receptor biogenesis. The Journal of biological chemistry 2009; 284: 14503–12.

48 Kawaguchi Y, Hagiwara D, Tsumura T, Miyata T, Kobayashi T, Sugiyama M, et al. Knockdown of endoplasmic reticulum chaperone BiP leads to the death of parvocellular AVP/CRH neurons in mice. Journal of neuroendocrinology 2023; 35: e13223.

49 Nakanishi T, Shimazawa M, Sugitani S, Kudo T, Imai S, Inokuchi Y, et al. Role of endoplasmic reticulum stress in light-induced photoreceptor degeneration in mice. Journal of neurochemistry 2013; 125: 111–24.

50 Oida Y, Hamanaka J, Hyakkoku K, Shimazawa M, Kudo T, Imaizumi K, et al. Post-treatment of a BiP inducer prevents cell death after middle cerebral artery occlusion in mice. Neuroscience letters 2010; 484: 43–6.

51 Inokuchi Y, Nakajima Y, Shimazawa M, Kurita T, Kubo M, Saito A, et al. Effect of an inducer of BiP, a molecular chaperone, on endoplasmic reticulum (ER) stress-induced retinal cell death. Investigative ophthalmology & visual science 2009; 50: 334–44.

52 Wang H, Lv S, Stroebel D, Zhang J, Pan Y, Huang X, et al. Gating mechanism and a modulatory niche of human GluN1-GluN2A NMDA receptors. Neuron 2021; 109: 2443–56 e5.

53 Yoshida H, Matsui T, Yamamoto A, Okada T, Mori K. XBP1 mRNA is induced by ATF6 and spliced by IRE1 in response to ER stress to produce a highly active transcription factor. Cell 2001; 107: 881–91.

54 Wang M, Cotter E, Wang YJ, Fu X, Whittsette AL, Lynch JW, et al. Pharmacological activation of ATF6 remodels the proteostasis network to rescue pathogenic GABA(A) receptors. Cell & bioscience 2022; 12: 48.

55 Di XJ, Han DY, Wang YJ, Chance MR, Mu TW. SAHA enhances Proteostasis of epilepsy-associated alpha1(A322D)beta2gamma2 GABA(A) receptors. Chemistry & biology 2013; 20: 1456–68.

