## Supplemental Figures for "Adapting the endoplasmic reticulum proteostasis rescues epilepsy-associated NMDA receptor variants"

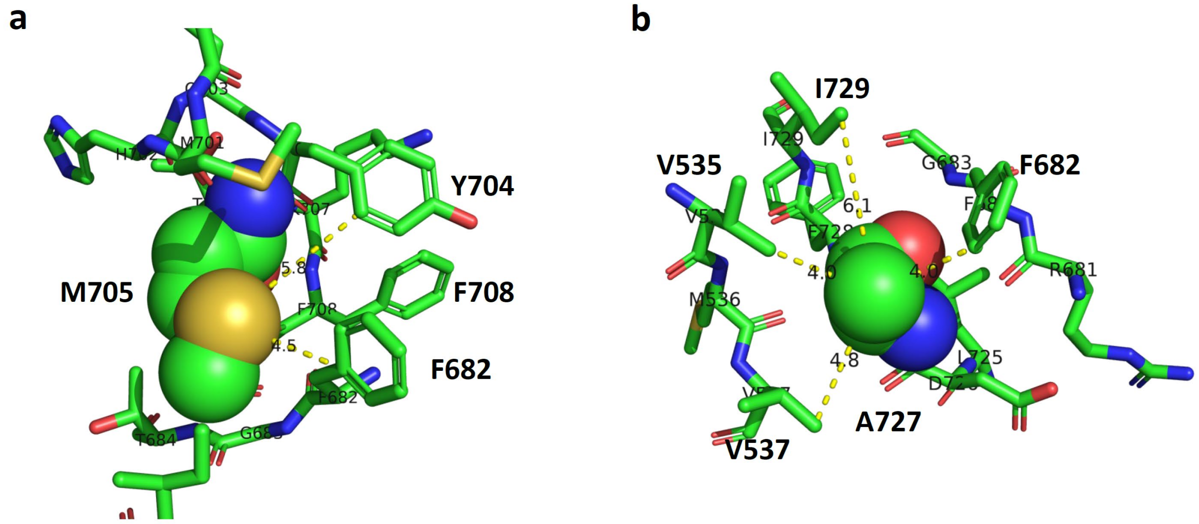


**Fig. S1. Vicinity of M705 and A727 in the NR2A subunit.** The NR2A residues are built from the cryo-EM structure of the human GluN1/GluN2A NMDA receptor in the glycine/glutamate bound state (PDB: 7EOS). **a**, M705 is shown as a sphere model and residues that are within 5 Å of M705 are shown as stick models. Aromatic residues, including F682, Y704, and F708, can form potential aromatic-Met interactions with M705. The closest distance between M705V sulfur atom and F682 is 4.5 Å and the closest distance between M705V sulfur atom and Y704 is 5.8 Å. **b**, A727 is shown as a sphere model and residues that are within 5 Å of A727 are shown as stick models. A727 can form potential hydrophobic interactions with other hydrophobic residues, including V535, V537, F682, A727, and I729.

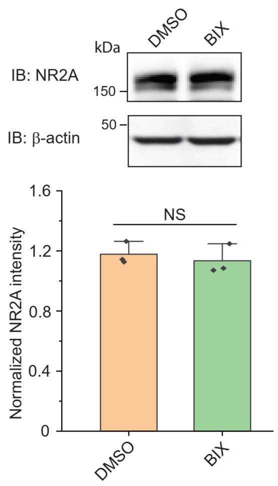


**Fig. S2. Effect of BIX on endogenous NR2A protein levels.** Western blotting analysis of the effect of BIX (10 μM, 24 h) on endogenous NR2A protein expression levels in human A2780 cells that express endogenous NMDA receptors. β-actin serves as the loading control of total protein lysates. Each data point was reported as mean ± SEM. Student's t-test (for comparison of two groups) was used for statistical analysis. NS: not significant.
